# Tying the Knot: Unraveling the Intricacies of the Coronavirus Frameshift Pseudoknot

**DOI:** 10.1101/2023.12.28.573501

**Authors:** Luke Trinity, Ulrike Stege, Hosna Jabbari

**Author notes:** Senior author (Ulrike Stege, Hosna Jabbari).

## Abstract

Understanding and targeting functional RNA structures towards treatment of coronavirus infection can help us to prepare for novel variants of SARS-CoV-2 (the virus causing COVID-19), and any other coronaviruses that could emerge via human-to-human transmission or potential zoonotic (inter-species) events. Leveraging the fact that all coronaviruses use a mechanism known as −1 programmed ribosomal frameshifting (−1 PRF) to replicate, we apply algorithms to predict the most energetically favourable secondary structures (each nucleotide involved in at most one pairing) that may be involved in regulating the −1 PRF event in coronaviruses, especially SARS-CoV-2. We compute previously unknown most stable structure predictions for the frameshift site of coronaviruses via hierarchical folding, a biologically motivated framework where initial non-crossing structure folds first, followed by subsequent, possibly crossing (pseudoknotted), structures. Using mutual information from 181 coronavirus sequences, in conjunction with the algorithm KnotAli, we compute secondary structure predictions for the frameshift site of different coronaviruses. We then utilize the Shapify algorithm to obtain most stable SARS-CoV-2 secondary structure predictions guided by frameshift sequence-specific and genome-wide experimental data. We build on our previous secondary structure investigation of the singular SARS-CoV-2 68 nt frameshift element sequence, by using Shapify to obtain predictions for 132 extended sequences and including covariation information. Previous investigations have not applied hierarchical folding to extended length SARS-CoV-2 frameshift sequences. By doing so, we simulate the effects of ribosome interaction with the frameshift site, providing insight to biological function. We contribute in-depth discussion to contextualize secondary structure dual-graph motifs for SARS-CoV-2, highlighting the energetic stability of the previously identified 3 8 motif alongside the known dominant 3 3 and 3 6 (native-type) −1 PRF structures. Integrating experimental data within minimum free energy (MFE) hierarchical folding algorithms provides novel structure predictions to distill the relationship between RNA structure and function. In particular, fully categorizing most stable secondary structure predictions via hierarchical folding supports our identification of motif transitions and critical site targets for future therapeutic research.

**Author summary:** Finding evolutionary connections between coronaviruses frameshift element RNA structures is a worthwhile goal in contributing to treatment development for afflicted human and animal populations. Predicting the most energetically favourable RNA secondary structures, and how they may form via the hierarchical folding hypothesis, is an efficient use of computational resources to shed light on RNA structure-function.

We used the KnotAli algorithm to obtain mutual information from 181 coronaviruses frameshift RNA sequences. Guided by this evolutionary information, we computed secondary structure predictions to allow comparison of marked similarities and subtle differences between SARS-CoV-2 and other coronaviruses frameshift element RNA structures. In addition, we applied the Shapify algorithm to predict secondary structures for extended SARS-CoV-2 frameshift element sequences informed by SHAPE reactivity data. Here we critically expand the known landscape of most stable −1 PRF secondary structure conformations, isolating the location of key secondary structure motif transitions that can improve site targeting of viral therapeutics. Our application of hierarchical folding algorithms contributes novel predictions of functional RNA structures, enhancing discussion of how secondary structures unfold or refold to regulate frameshifting in coronaviruses.

## 1 Introduction

The virus SARS-CoV-2 (genus *Betacoronavirus*, subgenus *Sarbecovirus* i.e., Severe acute respiratory syndrome Beta-coronavirus [1]) is responsible for the COVID-19 pandemic, leading to hundreds of millions of COVID-19 cases. Like all coronaviruses, SARS-CoV-2 viral RNA forms a functional structure referred to as a *pseudoknot* that, combined with a *slippery* RNA sequence, makes the ribosome prone to shifting into the −1 reading frame at the frameshift site [2–5]. Despite a high resolution of accuracy achieved for the study of specific RNA molecules, answering the question of how the frameshift event is regulated via the folding or unfolding of RNA structures remains elusive.

Given the high degree of complexity in predicting how the frameshift pseudoknot structure may be wedged or somehow possibly obstruct the entrance of the ribosome mRNA channel to initiate the frameshift event [6], we strive to fully understand the initial and subsequent folding of RNA molecules within a secondary structure model. To better inform and interpret tertiary structure experiments and simulations, our computational analysis explores the landscape of possibly pseudoknotted structures to elucidate key SARS-CoV-2 folding conformations. Experiments show that frameshift efficiency is significantly higher for an extended frameshift sequence (2924 nucleotides) than minimal frameshift sequence (92 nucleotides) [7]. Therefore, unfolding of longer range RNA structures must affect how the RNA refolds into a pseudoknot in proximity with the ribosome [8, 9]. Here we quest to further characterize the ensemble of most stable RNA secondary structures for extended frameshift sequences in SARS-CoV-2 and related coronaviruses.

Predicting and understanding frameshift inducing RNA structures in SARS-CoV-2 and related viruses is a critical target for therapeutic development [9, 10]. One example is intracellular small molecule therapy [6, 11–16], a strategy to limit viral fitness by disrupting the twisted RNA structure that contacts the ribosome at multiple locations [6]. Experiments demonstrate that specific compounds can inhibit frameshift efficiency in SARS-CoV-2 [6, 15] and other coronaviruses, affecting both humans and bats [17]. For example, KCB261770 was found to reduce frameshift efficiency in SARS-CoV-1, SARS-CoV-2, and MERS-CoV [18]. Indeed, novel viral genome targeting found the SARS-CoV-2 frameshift pseudoknot site to be the most effective location to limit viral reproduction [19].

Vigilance is necessary as coronaviruses continue to evolve in animal reservoirs. SARS-CoV-2 viral evolution analysis finds it most phylogenetically related to bat SARS-like coronaviruses [20]. An intelligent strategy to prepare for future SARS-CoV-2 variants or novel coronaviruses must leverage evolutionary structure information as well as the *hierarchical folding hypothesis*, in order to understand the role of initial and subsequent RNA folding within frameshift element mechanics, e.g., our earlier work in predicting secondary structures for the 68 nucleotide (nt) SARS-CoV-2 sequence [21]. Searching for commonality of structure features between coronaviruses contributes to broad spectrum pseudoknot therapeutic targeting, evidenced by molecular dynamic simulations of previously unknown 3-D structures for bat-coronavirus frameshift mechanics [22]. Previous analysis of length-dependent structures in different coronaviruses identified how sequences evolved to support a range of frameshift element structure motifs [9] (cf. Section 2.2).

Functional RNA structures that regulate the frameshift event have been studied for multiple viruses [23–25]. In particular, RNA sequences in the frameshift region have been observed to possess *conformational plasticity*, meaning they can form different configurations regulating frameshift efficiency [23]. For SARS-CoV-2, folding into multiple structures is a functional viral phenomenon evidenced by optical tweezers experiments [26, 27] as well as SHAPE-MaP (selective 2’-hydroxyl acylation analyzed by primer extension and mutational profiling) [28, 29] chemical probing of the SARS-CoV-2 viral genome *in vitro* [8] and *in vivo* [30–32]. Multiple unique stable conformations for the SARS-CoV-2 frameshift pseudoknot have been predicted and observed via various methods including crystallography, cryo-EM, and 3D-physics simulations [5, 8, 31, 33–35]. Thermal unfolding of RNA found major and minor paths from the folded to unfolded state, concluding that stability of transient (i.e., intermediate) states dictates folding paths [36].

Intense predictions efforts continue to build the structural model proposed for the exact SARS-CoV-2 −1 ribosomal frameshift mechanics. The RNA sequence itself is the primary factor in determining the structure of an RNA, ergo mutations to the frameshift element sequence can disrupt structure-function. Analyses of mutated SARS-CoV-2 RNA sequences and resulting structures find even single nucleotide mutations can substantially reduce frameshift efficiency [6, 14, 37]. Empirical evidence supports the specificity needed for the SARS-CoV-2 frameshift sequence, i.e., that mutations in this structural region are remarkably rare. The most prevalent mutations in the frameshift region (C13536U and C13378U) were each observed in only 0.12% of over 700, 000 sequences most recently recorded via GISAID at the time of this writing [38, 39]. Furthermore, there is some evidence of evolutionary convergence, with the most common mutation C13536U increasing sequence similarity between SARS-CoV-2 and MERS-CoV [40].

To contribute to comprehensive RNA structural knowledge we apply thermodynamic-based minimum free energy (MFE) algorithms to predict *crossing* RNA structures (cf. Section 2.1) that may regulate ribosome pausing mechanics in betacoronaviruses. Towards resilient SARS-CoV-2 therapeutic treatments, we substantially expand on our previous hierarchical folding (i.e., non-crossing RNA structure forms first [41–45]) investigation of the SARS-CoV-2 68 nt frameshift element sequence [21], by predicting secondary structures for extended length coronavirus frameshift sequences and incorporating sequence covariation information.

We utilize RNA *secondary structure* prediction algorithms, which output a set of base pairs where each base is in at most one pair. Each RNA loop (i.e., unpaired region closed by a base pair) is assigned an energy value. RNA structure abstraction, such as secondary structure, enables efficient computational modeling that provides structure information to help explain the role of RNA folding within important biological functions, such as protein synthesis. Predicting the tertiary (3D) structure of RNA [46–51] is crucial; however, it is more expensive computationally. 3D RNA models can be guided by secondary structure as an input constraint (for in-depth RNA tertiary structure prediction review, cf. Li et al. [52]).

To combat SARS-CoV-2 and prepare for novel coronavirus outbreaks, our strategy is to build and leverage coronaviruses structure-function information in support of frameshift-disrupting therapeutics. We focus on the secondary structure formation and function of the frameshift element viral RNA in betacoronaviruses, especially SARS-CoV-2. Specifically, we are interested in the most stable (i.e., MFE) initial stems, and subsequent hierarchical folding of viral RNA into possibly pseudoknotted secondary structures.

First, we sought to detect and utilize coronaviruses sequence covariation by applying *KnotAli* [53], a free energy minimization algorithm that merges conserved evolutionary information within a relaxed hierarchical folding approach. We utilized KnotAli to predict possibly pseudoknotted secondary structures for SARS-CoV-2 and related coronaviruses. We present and discuss our results for inter- and intra-coronavirus RNA structuredness. Predictions via KnotAli showcase evolved conformational flexibility based on detected covariation, and also how mutations change predicted structures for SARS-CoV-2 and related viruses, especially bat coronaviruses. We provide additional context for known covariation and associated base pairing [33] by predicting novel base pairs that possess strong covariation within the multiple sequence alignment.

Second, to explore SARS-CoV-2 frameshift pseudoknot motifs in longer sequences, we applied our hierarchical folding algorithm, *Shapify*, to predict possibly pseudoknotted secondary structures while incorporating reactivity data. Following the approach of Schlick et al. [33], which extended to a 222 nt coronavirus frameshift sequence element window, we obtained SARS-CoV-2 frameshift element structure predictions for sequences increasing in length from 90 nt to 222 nt using Shapify, in combination with genome wide *in vivo* and sequence specific *in vitro* SHAPE data. Our predictions allow comparison of energetic stability between known secondary structure motifs [33]. Predictions via Shapify unveil a diverse array of energetically favourable potential pseudoknots for the frameshift element that were previously unknown. Our SHAPE-informed structure prediction analysis includes detailed classification of critical pseudoknot motifs. We report complex pseudoknot predictions both upstream (5*^′^*) and downstream (3*^′^*) with respect to the *native* pseudoknot structure (cf. Section 2.2), including the traditional *attenuator* hairpin [54]. By extending SHAPE-informed hierarchical-folding structure prediction, our analysis more precisely describes the landscape of energetically favourable RNA structures, facilitating site-specific therapeutic targeting strategies.

## 2 Materials and methods

First, we introduce background information for RNA structure prediction and secondary structure motifs [21]. Second, we outline the hierarchical folding methods for secondary structure prediction. That is, we introduce KnotAli [53] for detection of covariation through alignment and secondary structure prediction informed by covariation; and the Shapify algorithm [21] for hierarchical folding with SHAPE data as soft-constraint.

### 2.1 RNA secondary structure prediction

Computational methods to predict RNA secondary structure identify nucleotide bases that form base pairs when RNA molecules fold. RNA folding refers to the process by which RNA acquires its structure through base pairing. Prediction of the secondary structure (set of all base pairs) for an RNA molecule is a more achievable intermediate task shedding light on ultimately desirable RNA tertiary structure prediction. Thus, we focus on the prediction of RNA secondary structure, where any RNA molecule is represented by its sequence *S* of length *n*.

Each RNA sequence has an alphabet of four bases: adenine (A), cytosine (C), guanine (G), and uracil (U). When an RNA structure forms, complementary bases pair together and form hydrogen bonds. *Canonical base pairs* are most stable pairings that occur when ‘A’ pairs with ‘U’, or if ‘G’ pairs with either ‘C’ or ‘U’. Each nucleotide base in the RNA sequence is referred to by its position in *S* indexed from 1 to *n* from 5*^′^* (left) to 3*^′^* (right) end. A *base pair* is defined as the pairing of two bases *i* and *j* where 1 *≤ i < j ≤ n*, and represented as *i.j*. Each base can pair with at most one other base (i.e., no base triplets are allowed, which is left to tertiary structure prediction). Consecutive base pairs are referred to as a *stem*.

The free energy of an RNA structure is calculated as the sum of the energies of its loops, i.e., hairpin, bulge, internal loop or multiloop. For this, some loop energies were experimentally determined, where experimental results were not available, loop energies are extrapolated [55–57]. Base pairs *i.j* and *i^′^.j^′^* are *nested* if 1 *≤ i < i^′^ < j^′^ < j ≤ n*, and *disjoint* if 1 *≤ i < j < i^′^ < j^′^ ≤ n*. An RNA structure with only nested and disjoint base pairs is referred to as a *pseudoknot-free* structure. An RNA structure is considered *pseudoknotted* when at least two of its base pairs, *i.j* and *i^′^.j^′^* cross: 1 *≤ i < i^′^ < j < j^′^ ≤ n*, in which case both *i.j* and *i^′^.j^′^* are considered pseudoknotted base pairs.

In this work we assume RNA folds sequentially, supported by the *hierarchical folding hypothesis* [41]: *initial* stems first fold into a pseudoknot-free structure. Subsequently, additional base pairing can lower the free energy of the structure and possibly form pseudoknots. Hierarchical folding paths were experimentally identified in multiple pseudoknots [44], including frameshifting pseudoknots [45]. Computational methods for prediction of RNA secondary structure based on the hierarchical folding hypothesis, including KnotAli [53] and Shapify [21], find the MFE (most stable) structure for a given sequence *S*, and a pseudoknot-free input structure *G*.

RNA structure SHAPE chemistry experiments [42] and computational RNA folding simulations [43] find that initial structures may be modified locally to accommodate formation of more stable base pairs. Therefore, the KnotAli and Shapify algorithms are equipped to allow for minor modification to improve their prediction accuracy.

### 2.2 SARS-CoV-2 frameshift secondary structure motifs

SARS-CoV-2 secondary structure prediction efforts find multiple length-dependent structural *motifs* related to the frameshift event [9, 33, 58]. We use *dual graph* nomenclature introduced in [33, 59–61] to refer to the RNA secondary structure or substructure predicted at or directly 3*^′^* of the coronavirus frameshift element slippery sequence. A dual graph specifies connectivity and topological aspects of secondary structure by representing each stem by a vertex (cf. Fig 1). Dual graph edges between vertices represent junction of stems, or loops. Specifically, each unpaired RNA strand is represented by an edge.

**Fig 1.**
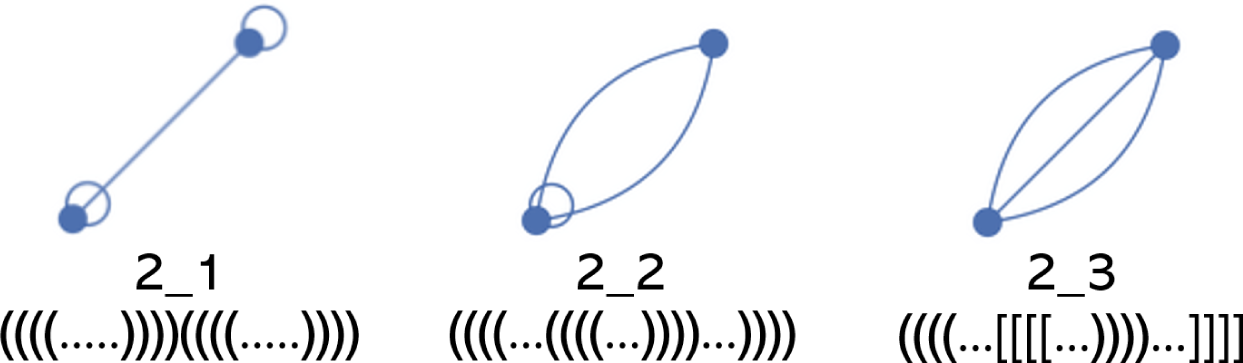
RNA dual graph motifs and nomenclature with two vertices. Vertices represent stems. Edges represent junction of stems, or bulge/internal loops with more than one residue on each strand. Self-edges represent hairpin loops. Dual graphs are referred to by two numbers, listed below each respective graph. The first number indicates the number of vertices, the second number specifies the topology, e.g., 2 1 is the dual graph secondary structure motif with two vertices, specifically the first possible topology. For additional details refer to the RNA-As-Graphs database [59]. Dot bracket example structures for each respective motif shown below number labels. Open parentheses/brackets show the base on the 5*^′^* side of the sequence, closed parentheses/brackets represent the base on the 3*^′^* side of the sequence that are binding together. Each period “.” represents an unpaired base.

Dual graph representations allow for variable length of stems and loops, which makes RNA secondary structure pattern or motif identification easier. Note that the number of possible topologies rises exponentially with the number of stems, e.g., with three stems, eight possible topologies, with six stems, 508 possible topologies. Therefore, we use dot bracket notation (cf. Fig 1) or arc diagrams (cf. Fig 2) to represent secondary structure predictions with more than three stems, such as those predicted for the SARS-CoV-2 sequence of length 222 which may have ten or more stems. In arc diagrams, the RNA sequence of bases is shown from left to right (5*^′^* to 3*^′^*) in a single horizontal line. Base pairs are represented as arcs between bases.

**Fig 2.**
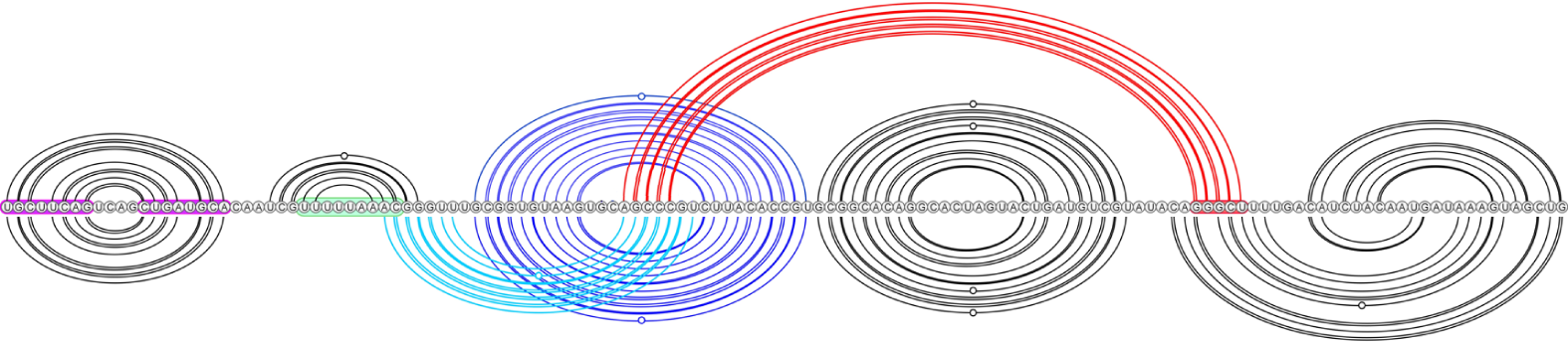
Dominant SARS-CoV-2 pseudoknot motif predictions via Shapify. SARS-CoV-2 frameshift element sequence shown as a horizontal line from 5*^′^* (left) to 3*^′^* (right). Arcs represent predicted base pairs. Top arc diagram includes 3 6 motif component (cf. Table 2 for dot-bracket format) of the fifth most stable structure predicted via Shapify (cf. Sections 2.4-2.6) for 144 nt sequence, free energy −29.45 kcal/mol, initial stem 5 base pairs in red (free energy −4.22 kcal/mol). Downstream pseudoknot target sequence highlighted in red. Bottom arc diagram includes 3 3 motif component of the MFE structure predicted via Shapify for 144 nt sequence, free energy −30.93 kcal/mol, initial stem 2 base pairs in light blue (free energy −6.1 kcal/mol). Initial stem 1 base pairs in dark blue (free energy −11.48 kcal/mol).

The *native* frameshift element structure for SARS-CoV-2 [54], also referred to as 3 6 motif [33], is a three-stemmed pseudoknot forming directly downstream of the slippery RNA sequence (cf. Fig 2 top arc diagram, pseudoknotted base pairs in red). Structure-function research also finds a frameshift efficiency downregulation mechanism via a simple RNA loop preceding the slippery sequence, i.e., the *attenuator* hairpin, via interaction with the ribosome during translocation or elongation [27, 54] (cf. Fig 2, sequence highlighted in pink). Previously, we predicted 3 6 motif structures and structure similarity with SARS-CoV-2 for SARS-CoV-1 and MERS-CoV [21].

Within a hierarchical folding framework for the SARS-CoV-2 frameshift element sequence, the stem occurring directly downstream of the slippery sequence, referred to as stem 1, was identified as the most energetically favourable initial stem (cf. Fig 2, dark blue arcs) within a 68 nt window [21]. Likely due to its high relative stability within the energy landscape, stem 1 refolds quickly when unfolded [26]. Stem 1 was also found to be highly conserved among coronaviruses [33], and present in both major structure motifs proposed by Schlick et al. [33] (3 6 for 77 nt window, and 3 3 for 144 nt window). For extended sequence lengths, native stem 1 may not form, instead, upstream base pairing has been identified [7].

Conversely, the second most energetically favourable initial stem for SARS-CoV-2 frameshift element sequence, i.e., stem 2, was found to pair differently depending on window size around the frameshift site considered for prediction. Schlick et al. [33] used secondary structure prediction in combination with SHAPE structural probing and thermodynamic ensemble modeling to identify stem 2 either (*A*): paired into a simple pseudoknot crossing upstream of stem 1 in the 3 3 motif (95.6% of ensemble at 144 nt, cf. Fig 2, light blue arcs), or (*B*): paired downstream forming the native H-type pseudoknot in the 3 6 motif (98% of ensemble at 77 nt).

### 2.3 Covariation-informed hierarchical folding

KnotAli [53] combines the strengths of MFE prediction and alignment-based methods through relaxed hierarchical folding. KnotAli uses a multiple RNA sequence alignment as input to predict possibly pseudoknotted secondary structures for each sequence in the alignment. KnotAli first identifies a set of pseudoknot-free base pairs, based on mutual information, to guide subsequent free energy minimization. Note that relaxed hierarchical folding indicates predictions can be reached via multiple different paths allowing suboptimal structures, and the initial structure may be modified to accommodate base pairs that lower the free energy of the structure. Output from KnotAli includes base pairs that show strong covariation among the multiple sequence alignment, as well as possibly pseudoknotted predictions for each individual sequence.

### 2.4 SHAPE-informed hierarchical folding

Shapify [21] is a relaxed hierarchical folding algorithm that uses as input an RNA sequence, a pseudoknot-free structure, and a SHAPE reactivity dataset, to predict a possibly pseudoknotted secondary structure. Shapify computes a secondary structure prediction via four different biologically supported methods and returns the MFE structure from the four methods. With respect to the four structures from the four methods, we include in our results any additional *suboptimal* structures that fall within 2 kcal/mol of the MFE prediction.

To create the input structure following the hierarchical folding hypothesis, we used the HotSpots package (cf. HotKnots V2.0 [57]) and identified up to 20 lowest free energy unique stems (referred to as *initial stems*) for each respective window to be used as constraint via Shapify. The stems were ranked from most stable to least stable based on their free energy and are referred to by their ranking as their IDs.

### 2.5 Data

We obtained the coronavirus alignment of Schlick et al. [33], where out of 3760 SARS-CoV-2 coronavirus sequences [38], and 2855 other coronavirus sequences [62], 1248 sequences were found to be non-redundant [63]. These 1248 sequences were structurally aligned to the 222 nt SARS-CoV-2 frameshift element SHAPE consensus structure [33] using the Infernal covariance model [64] giving a final result of 182 non-duplicate homologous sites including seven SARS-CoV-2 sequences. Here, we converted the alignment of 182 sequences to FASTA format for input into KnotAli.

For input to Shapify we utilized the reference genome for SARS-CoV-2 from the National Center for Biotechnology Information, *NC* 045512.2 [65]. We obtained three available SARS-CoV-2 SHAPE datasets to guide Shapify predictions, two *in vivo*: Huston et al. [30], and Yang et al. [32], one *in vitro*: Manfredonia et al. [31].

### 2.6 Shapify window procedure

Secondary structure predictions were obtained via Shapify with initial stems and SHAPE data for each SARS-CoV-2 sequence varying in length from 90 nt to 222 nt, with a step size of 1. Therefore, our results are based on removing successive nucleotides from the 5*^′^* side of the RNA sequence for a total of 132 sequences analyzed (cf. Table 1 for window position and length). To compare free energy of structures for different sequence lengths, we divided the energy of each structure by its respective sequence length for an unbiased inter-window comparison, referred to as free energy per nt (for visualization see Fig 6): Free energy per nt = Structure free energy Sequence length

**Table 1.** Shortest and longest window sizes used for SARS-CoV-2 structure predictions via Shapify.

| Position | Length |
| --- | --- |
| 13485 – 13575 | 90 |
| 13354 – 13575 | 222 |

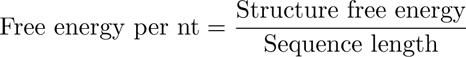

## 3 Results

We first visualize base pairs with strong covariation among the multiple sequence alignment identified by KnotAli. Next, we present secondary structure predictions for coronaviruses frameshift element sequences via KnotAli. Then, we display our Shapify structure investigation applying hierarchical folding to predict SARS-CoV-2 secondary structures for frameshift element sequences up to length 222. All structure predictions and associated data can be found in S1 File.

### 3.1 KnotAli secondary structure predictions

KnotAli finds strong covariation in the multiple sequence alignment for base pairs that align with the known attenuator hairpin [54], and also the native 3 6 motif (cf. Fig 3).

**Fig 3.**
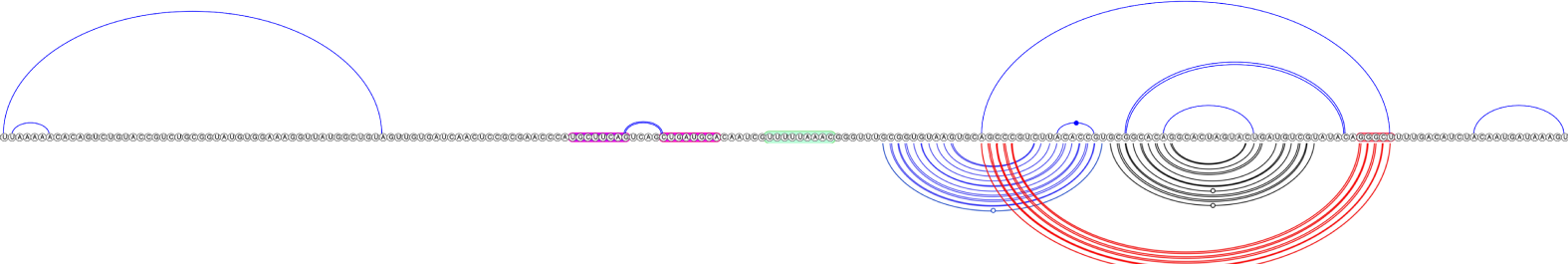
Coronavirus frameshift element covariation. Base pairs in the top arc diagram have strong covariation among the multiple sequence alignment identified by KnotAli. Bottom arc diagram displays the SARS-CoV-2 native 3 6 psuedoknot, downstream target sequence in red. SARS-CoV-2 attenuator hairpin sequence highlighted in pink, slippery sequence in green.

Among the seven SARS-CoV-2 sequences in the alignment, structure predictions for five sequences included the 3 3 motif, while two sequences (EPI ISL 465643, EPI ISL 426088) resulted in a prediction that included the native 3 6 motif instead (cf. Fig 4). A structure containing the 3 6 motif was also predicted for BtRf-BetaCoV (cf. Fig 5). BtRf-BetaCoV has 77% overall identity with SARS-CoV-2, and 91% frameshift sequence identity (FSID) with SARS-CoV-2 as identified by Schlick et al. [33].

**Fig 4.**
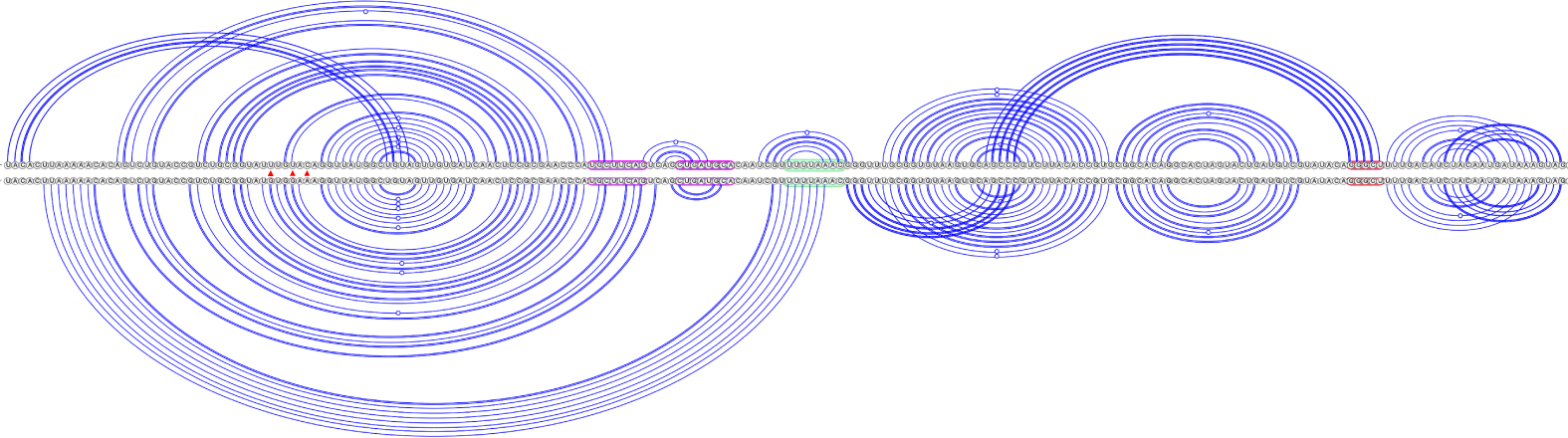
SARS-CoV-2 secondary structure predictions via KnotAli. Top arc diagram: free energy −36.47 kcal/mol, EPI ISL 426088, includes 3 6 motif. Bottom arc diagram: free energy −40.65 kcal/mol, EPI ISL 426905, includes 3 3 motif. Mutations indicated with a red *△* symbol between sequences. Attenuator hairpin sequence highlighted in pink, slippery sequence in green, downstream native pseudoknot target sequence in red.

**Fig 5.**
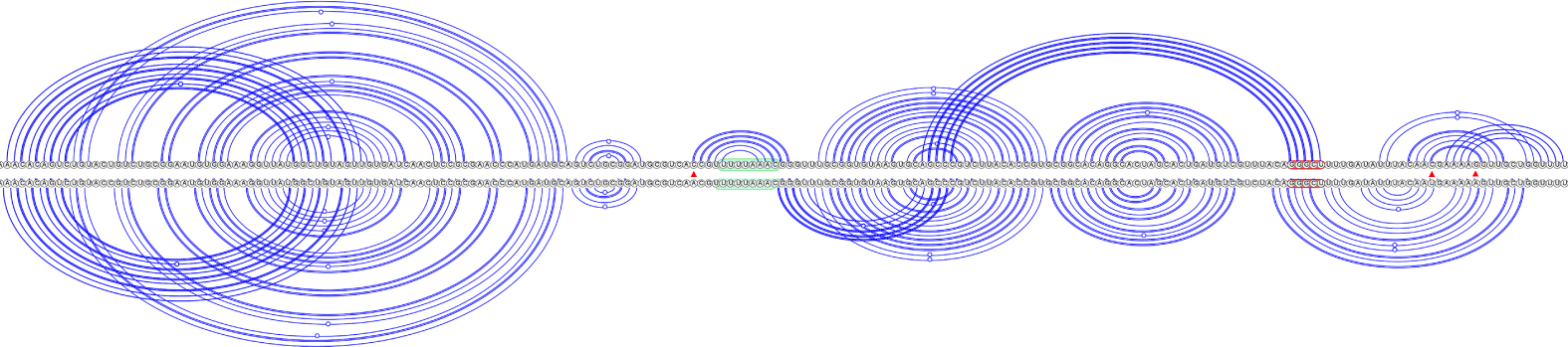
Bat coronaviruses secondary structure predictions via KnotAli. Top arc diagram: BtRf-BetaCov, free energy −43.24 kcal/mol, KJ473811, includes 3 6 motif. Bottom arc diagram: SARS-like WIV1-CoV, free energy −39.72 kcal/mol, KU444582, includes 3 3 motif. Mutations indicated with a red *△* symbol between sequences. Slippery sequence in green, downstream native pseudoknot target sequence in red.

The majority of sarbecovirus secondary structure predictions include the 3 3 motif: Pangolin-CoV (98% FSID), SARS-CoV-1 (93% FSID), SARS-like WIV1-CoV (93% FSID, cf. Fig 5), BtRs-BetaCoV (92% FSID), Bat-Cov-Cp (91% FSID), and Bat-CoV-Rp (91% FSID).

Sarbecovirus predictions demonstrate significant structuredness. Pseudoknots were predicted upstream of the native frameshift pseudoknot site for SARS-CoV-2, SARS-CoV-1, SARS-like WIV1-CoV, BtRs-BetaCoV, BtRf-BetaCoV, and Bat-CoV-Cp. Pseudoknots were predicted downstream of the native frameshift pseudoknot site for Pangolin-CoV, SARS-like WIV1-CoV, BtRf-BetaCoV, and Bat-HpBetaCoV.

### 3.2 Shapify secondary structure predictions

Following the procedure outline in Section 2.6, we obtained a total of 10, 916 secondary structure predictions via Shapify (cf. Fig 6) for the SARS-CoV-2 frameshift element window of varying length. We observe that the most stable secondary substructures (including only up to three stems at or directly 3*^′^*of the slippery sequence) can be classified into four pseudoknot dual-graph motifs: 2 3, 3 3, 3 6 (native-type), and 3 8 (cf. Fig 2, Fig 6, and Table 2).

**Fig 6.**
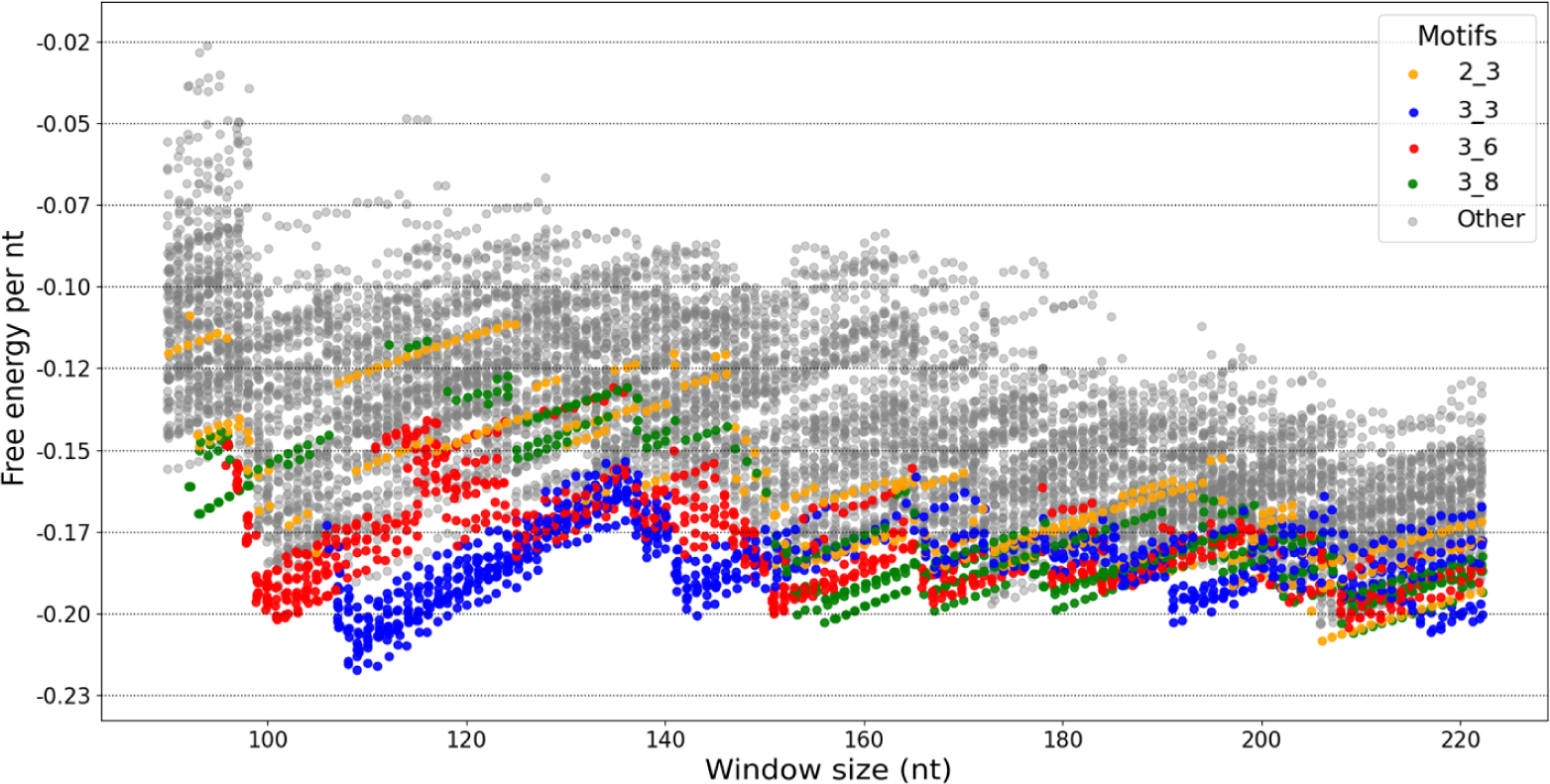
SARS-CoV-2 secondary structure motifs free energy per nt. Each point represents a Shapify predicted secondary structure for the SARS-CoV-2 frameshift sequence (cf. Section 2.6, Table 1). X-axis represents window size, y-axis represents free energy per nt. Dots are colored based on the four listed dual-graph motifs (legend in top-right) detected at or directly 3*^′^* of the slippery sequence (cf. Table 2), or grey for none.

**Table 2.**
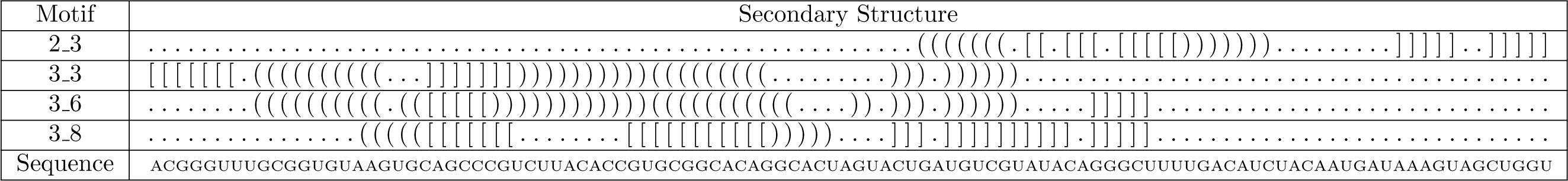
Secondary structure pseudoknot motifs in dot bracket notation. Note that motif classification allows minimal modification to structures, e.g., in the size of loops. Sequence ID: *NC* 045512.2 [65], indices 13467 − 13565. Open parentheses/brackets show the base on the 5*^′^*side of the sequence, closed parentheses/brackets represent the base on the 3*^′^* side of the sequence that are binding together. Each period “.” represents an unpaired base.

Our predictions confirm that initial stem 1 folds into the 3 3 motif, as part of the complete MFE structure, for three specific window size intervals: 107 − 151, 191 − 199, and 216 − 222 (cf. Fig 6). There was path convergence, meaning multiple initial stems resulted in different predictions that each contain the 3 3 motif (cf. Fig 7). For example, with the 144 nt sequence as input, the MFE structure and the next four most stable structures all include the 3 3 motif. At the critical window sizes 98 − 106, which include the region directly preceding the slippery sequence, the 3 3 motif is sufficiently destabilized leading to the MFE structure including the 3 6 motif instead. With window sizes 152 − 153, the MFE structure again includes the 3 6 motif as a key component. At the shortest length, as the 3 6 motif is destabilized, a 3 8 motif is predicted within the MFE structure for window sizes 93 − 97.

**Fig 7.**
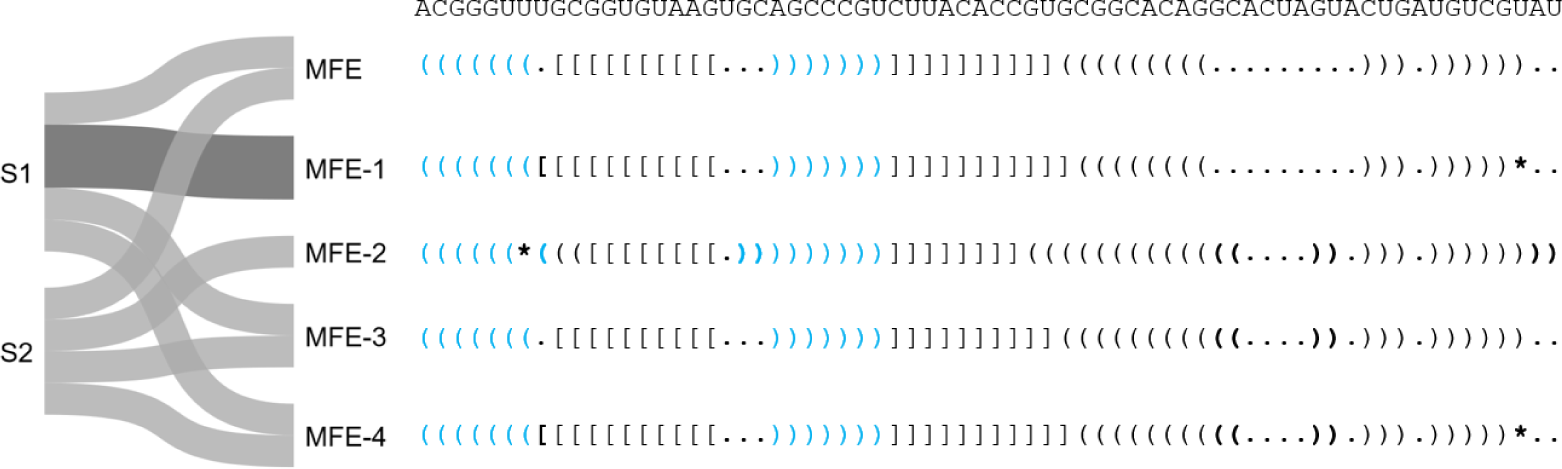
Convergence to the most stable structures that contain the 3 3 motif. 144 nt window Shapify predictions with initial stems 1 and 2 as constraint result in the MFE and most stable structures that contain the 3 3 motif. Initial stems labeled on the left (e.g., S1 for initial stem 1). Darker grey path indicates the structure predicted with a specific initial stem was the same for two SHAPE datasets. Light grey path indicates the structure predicted with an initial stem was specific to one SHAPE dataset. Structures on the right are labeled by energy proximity to the MFE structure, e.g. MFE-1 is the lowest free energy structure after the MFE structure. Differences from the MFE structure are marked in bold, with parenthesis representing changes in paired bases, and asterisks representing predicted unpaired bases that were paired in the MFE structure. 3 3 motif pseudoknotted base pairs shown in light blue.

Predictions via Shapify unveil changes in energetic favourability of pseudoknotted motifs for extended length sequences. We find the 3 8 motif predicted within the MFE structure for window sizes of 154 − 172 and 179 − 185. In addition, a 2 3 motif is predicted within the MFE structure for window sizes 205 − 208. The 2 3 motif is energetically close to the 3 8 motif at window sizes 209 − 215. Beyond local pseudoknotted motifs, we detect additional stable pseudoknotted regions suggesting possible upstream and downstream structure-function of the frameshift element (cf. Fig 8,9 and 10).

**Fig 8.**
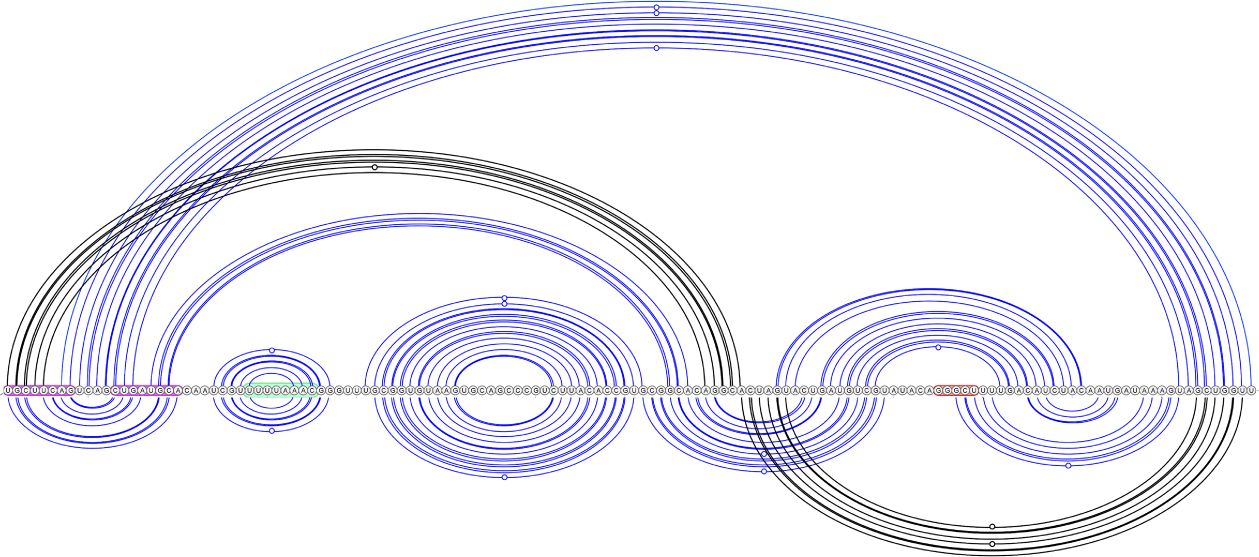
Structural regions involving pseudoknots in SARS-CoV-2, 144 nt window via Shapify. Top arc diagram: MFE-20, free energy –24.29 kcal/mol, initial stem 18 in black (free energy –0.35 kcal/mol). Bottom arc diagram: MFE-12, free energy –25.76 kcal/mol, initial stem 11 in black (free energy –1.44 kcal/mol). Attenuator hairpin sequence in pink, slippery sequence in green, downstream native pseudoknot pairing region in red.

**Fig 9.**
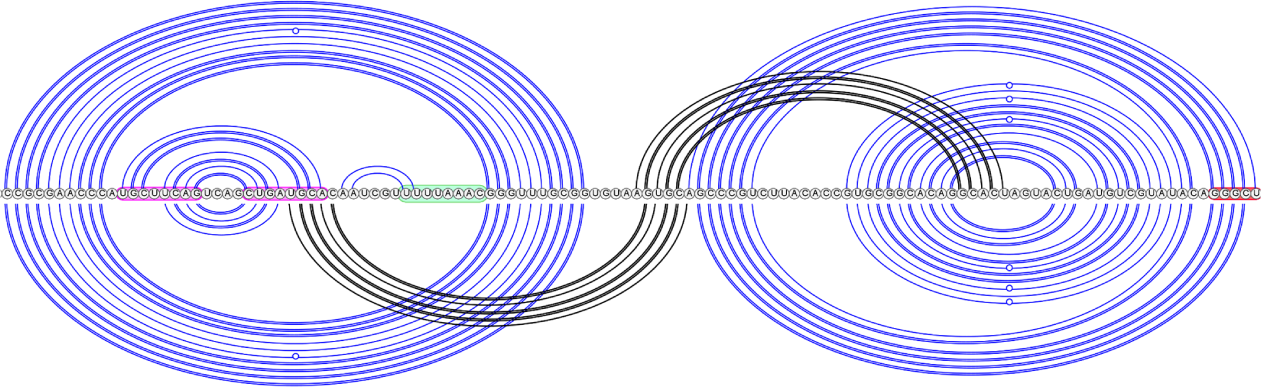
SARS-CoV-2 pseudoknot predictions overlap, 222 nt window via Shapify. Top arc diagram: MFE-5, free energy –45.04 kcal/mol, initial stem 15 in black (free energy –2.74 kcal/mol). Note that the MFE-5 pseudoknot was also detected within the 144 nt window (cf. MFE-19) and the 68 nt window [21]. Bottom arc diagram: MFE-29, free energy –39.34 kcal/mol, initial stem 12 in black (free energy –3.41 kcal/mol). Attenuator hairpin sequence in pink, slippery sequence in green, downstream native pseudoknot pairing region in red.

**Fig 10.**
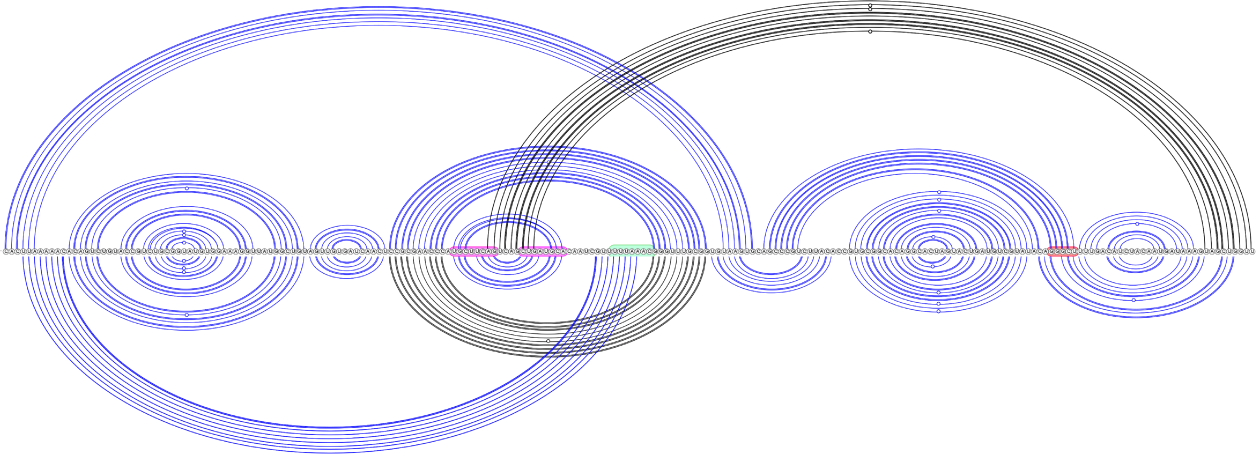
SARS-CoV-2 long range pseudoknot predictions, 222 nt window via Shapify. Top arc diagram: MFE-58, free energy –34.16 kcal/mol, initial stem 16 in black (free energy –1.98 kcal/mol). Bottom arc diagram: MFE-10, free energy –42.74 kcal/mol, initial stem 2 in black (free energy –10.08 kcal/mol). Attenuator hairpin sequence in pink, slippery sequence in green, downstream native pseudoknot pairing region in red.

## 4 Discussion

Fully understanding functional RNA structures remains an elusive but worthwhile goal, and is a necessary step towards effective therapeutics for viral infections. Despite many attempts to distill how coronavirus RNA folds to efficiently regulate a frameshift event, Structures on the right are labeled by energy proximity to the MFE structure, e.g. MFE-1 is the lowest free energy structure after the MFE structure. Differences from the MFE structure are marked in bold, with parenthesis representing changes in paired bases, and asterisks representing predicted unpaired bases that were paired in the MFE structure. 3 3 motif pseudoknotted base pairs shown in light blue. much is still unknown about the mechanism. We employed two hierarchical folding free energy minimization algorithms, KnotAli, and Shapify, for prediction of possibly pseudoknotted structures of SARS-CoV-2. Next, we discuss and contextualize our predictions of the coronavirus frameshift element to advance known structure information towards site-specific viral therapeutics, namely: (1) secondary structure predictions for SARS-CoV-2 and bat coronaviruses frameshift elements via KnotAli, and (2) insights from SHAPE-informed hierarchical folding predictions for SARS-CoV-2 via Shapify.

### 4.1 Coronaviruses frameshift element covariation

With regard to the base pairs detected by KnotAli that show strong covariation among the multiple sequence alignment, we first note the innermost base pair of the traditional attenuator hairpin stem-loop (GC base pair at 13442.13447). Covariation of this innermost base pair, along with the attenuator hairpin being functionally retained in SARS-CoV-1 and SARS-CoV-2 despite sequence differences [54], suggests the loop may be structurally conserved. Compensatory mutations related to the GC base pair at 13442.13447 include a potential AU base pair at 13442.13447 in transmissible gastroenteritis virus (TGEV, 59% FSID), porcine respiratory coronavirus (PRCV, 60% FSID), and turkey coronavirus (Turkey-CoV, 60% FSID), also a potential UA base pair at 13442.13447 in Night heron coronavirus (Night-Heron-CoV, 58% FSID). In addition, the outermost base pair of the pseudoknotted stem in the native 3 6 motif structure is found to have strong covariation, supporting previous results of Schlick et al. [33].

A two-branched multiloop directly downstream of the slippery sequence is identified as having strong covariation by KnotAli. The larger branch forms a bulge loop via a GC base pair at 13507.13536. Covariation of the GC base pair at 13507.13536 is supported by possible GU base pair at 13507.13536 in Pangolin-CoV, and also possible AU base pair at 13507.13536 in TGEV, Night-Heron-CoV, PRCV, Feline-CoV (61% FSID), and Canine-CoV (58% FSID). Further support is seen via possible UG base pair at 13507.13536 in infectious bronchitis virus (IBV, 60% FSID) and Turkey-CoV. Covariation of this bulge loop inner base pair, AU at 13512.13524, is supported by possible UA base pair at 13512.13524 in TGEV, PRCV, Feline-Cov, Canine-CoV and BtSk-Alpha-CoV (61% FSID), and also possible GU base pair at 13512.13524 in IBV and Turkey-CoV. In addition, the AU base pair at 13512.13524 is predicted via Shapify in multiple different SARS-CoV-2 pseudoknots including the 3 3, 3 6, and 3 8 motifs (cf. Table 2) and longer range predictions (cf. Fig 9 and 10).

With respect to the native frameshift pseudoknot (cf. Fig 3), there are two base pairs with strong covariation predicted upstream, and one base pair with strong covariation predicted downstream. Previously, Schlick et al. [33] identified an upstream AU base pair at 13366.13411 in SARS-CoV-2, Pangolin-CoV, Bat-CoV-Rp, BtRs-BetaCoV, SARS-like WIV-1-CoV, SARS-CoV, BtRf-BetaCoV, and Bat-CoV-Cp. Their covariance analysis highlighted compensatory mutations via a CG base pair at 13366.13411 in TGEV and PRCV, a UG base pair at 13366.13411 in Feline-CoV, and a UA base pair at 13366.13411 in Rousettus-Bat-Cov (64% FSID). KnotAli finds strong covariation in the multiple sequence alignment to support a UA base pair at 13360.13366, with A13366 pairing upstream with U13360 as opposed to the previously identified AU base pair at 13366.13411. This UA base pair at 13360.13366 is supported by compensatory mutations via possible GU base pair at 13360.13366 in TGEV, PRCV, Feline-CoV and Canine-Cov. Finally, the additional upstream base pair predicted via KnotAli, UA at 13359.13410, is supported by possible GC base pair at 13359.13410 in IBV, Rousettus Bat-CoV, and Turkey-CoV. KnotAli also finds strong covariation in a downstream AU base pair at 13553.13565, supported by possible GC base pair at 13553.13565 in IBV and Turkey-CoV.

These novel predicted base pairings, supported by coronavirus multiple sequence alignment covariation, provide additional context to previously identified covariation in coronaviruses [33]. As a constraint guiding secondary structure prediction via KnotAli, we note the covariation-informed initial structure is preserved in certain individual predictions for coronaviruses, but in other cases more stable structures can be reached by disrupting these base pair(s). We observe that minimal upstream mutations to SARS-CoV-2 sequences lead to either the 3 3 motif or 3 6 motif predicted via KnotAli (cf. Fig 4). In addition, minor changes between bat coronaviruses like BtRf-BetaCov and SARS-like WIV1-Cov not only led to different motif predictions, but also affect predicted downstream pseudoknot pairings (cf. Fig 5). We conclude that minimal upstream and downstream sequence variation can significantly change conformations of the frameshift element. In supporting resilient frameshift therapeutics, it could be valuable to assess the viability of treatments intended for SARS-CoV-2 across various bat coronaviruses sharing similar sequences and structures. Such investigations could provide more detailed insights into how the frameshift regulation mechanism may be disrupted.

### 4.2 SHAPE-informed hierarchical folding

Our secondary structure predictions for the SARS-CoV-2 frameshift sequence via Shapify offer insight into frameshift dynamics. By considering a SARS-CoV-2 sequence of varying length for prediction, we simulated how the interaction of the ribosome with the frameshift element [2] affects RNA structural motifs of SARS-CoV-2. In doing so we have characterized the landscape of possibly pseudoknotted structures, finding the 3 8 motif to be the most energetically favourable at specific sequence lengths, while also clarifying interplay between the dominant 3 3 and 3 6 native-type −1 PRF structures.

Irrespective of length, structures containing the 3 3 motif reached the lowest free energy per nt (nearly −0.23, cf. Fig 6). Our SHAPE-informed hierarchical folding also demonstrates innate resiliency, i.e., redundant folding paths from different initial stems all leading to the 3 3 motif structure. This extends previous results finding similar path convergence to the 3 6 motif structure within a smaller 68 nt window [21].

Secondary structure predictions demonstrate that as shorter sequence length destabilizes the 3 3 motif, the 3 6 native-type motif seamlessly emerges to dominate the ensemble, suggesting a transition between the two motifs. We observe the shift from 3 3 to 3 6 as occurring at SARS-CoV-2 sequence index 13468, when the second of three successive guanine nucleotides that would otherwise form the 3 3 motif stem is removed from the sequence under consideration. This location for a potential secondary structure transition was also identified within a partition function framework for the SARS-CoV-2 frameshift element [66]. Beyond these two major motifs, additional pseudoknot patterns are needed to fully describe the MFE structures of the ensemble, especially the 2 3 and 3 8 motifs.

The 3 8 motif has been previously detected as part of the SARS-CoV-2 frameshift element by *in vivo* probing and subsequent folding analysis within a 126 nt window [30]. It was also predicted computationally via hierarchical folding informed by SHAPE data as soft-constraint within a 68 nt window [21]. Observing that the 3 8 motif is the most energetically favourable structure for shorter length sequences, we hypothesize it may act as a transient structure facilitating refolding of either the 3 3 or 3 6 motif. In further support of a link between these three motifs is that they share an adenine bulge (A13524), which has long been identified as critical in frameshift regulation [67].

Further structure analysis is needed to understand function of the 2 3 motif, which is known to be prevalent in viral frameshift elements, especially plant viruses, but also simian retrovirus type-1, and mouse mammary tumor virus [61]. We hypothesize this downstream pseudoknot folding may have some effect on translation termination of the ribosome, which has been linked to frameshift regulation [68].

Our novel predictions suggest function of the attenuator hairpin via previously unknown pseudoknotted base pairing, paving the way for future tertiary modeling of SARS-CoV-2 frameshift RNA structure-function. Attenuator hairpin alternative pairings which include the slippery sequence and initial stem 1 have also been proposed [7]. We predict for the first time attenuator hairpin bases folding into long range pseudoknotted interactions (cf. Fig 8, top arc diagram, and Fig 10, top arc diagram partially supported by IPKnot structure prediction [33, 69]), and also a potential H-type pseudoknot structure with the stem-loop conserved as part of a pseudoknotted multiloop (Fig 9, bottom arc diagram). Notably, we visualized multiple structures that possess significant overlap or regions of structural similarity. While certain stems overlap, other pseudoknotted base pairs form which indicate potential conformational switching between the two structures.

The latest SARS-CoV-2 tertiary structure experimental results required molecular simulation of additional folding to reach higher agreement with previous experimentally derived molecular structures [35]. Understanding how initial most stable stems give way to pseudoknotted motifs via unfolding and refolding may unlock the key to discerning kinetic trajectories, which are currently not well defined. With regard to the most stable initial stems obtained via HotKnots [70], in general we find initial stem 1, i.e, stem 1 (free energy −11.48 kcal/mol, cf. Fig 2, dark blue base pairs) is the lowest free energy stem. This confirms previous results finding initial stem 1 to be most stable within a 68 nt window [21]. Other initial stems can change in their relative energetic stability, e.g., initial stem 2 (free energy −6.1 kcal/mol, cf. Fig 2, light blue base pairs) is the second lowest free energy stem for the 144 nt window, but this is supplanted in the 222 nt window by an additional highly stable upstream stem. We hypothesize a possible frameshift downregulation function for this additional highly stable stem, because in our analysis no folding path was detected to the most stable secondary structures. Overall, further study is needed to fully understand the mechanics of the folding from initial stems into secondary structures, especially in pseudoknots.

In the folding process of the SARS-CoV-2 frameshift pseudoknot, the native pseudoknotted stem likely folds last [26]. Hence, we suggest exploring site-specific therapeutic targeting of the downstream native pseudoknot pairing. Since this pseudoknot critically refolds to initiate the frameshift, it should be an accessible location to disrupt the frameshift pseudoknot. In comparing the site target potential of an overlapping 5*^′^* site that did not include the pseudoknot target, oligonucleotide targeting found the downstream native pseudoknot pairing to be a more effective target for reducing frameshift efficiency [71]. The downstream native pseudoknot site may be even more effective than previously realized, because it includes key structure pairings of the 3 8 and 2 3 motifs, which are each predicted within the MFE structure at specific sequence lengths. For more comprehensive and broadly applicable therapeutics, upstream sites, like the attenuator hairpin, should also be considered in subsequent work. Further exploration of the intricate relationship between RNA structure and function is needed in the field of coronavirus therapeutics, towards more positive outcomes for human and animal health.

## Supporting information

**S1 File. Supplementary Materials.**

## Acknowledgments

We thank and acknowledge the Computational Biology Research and Analytics Lab for invaluable feedback, and the Microsoft AI for Health Azure Grant for funding support.

